# Taxonomic and functional diversity of aquatic heterotrophs is sustained by dissolved organic matter chemodiversity

**DOI:** 10.1101/2022.03.21.485019

**Authors:** Sarahi L. Garcia, Julia K. Nuy, Maliheh Mehrshad, Justyna J. Hampel, Vicente T. Sedano-Nuñez, Moritz Buck, Anna-Maria Divne, Eva S. Lindström, Daniel Petras, Jeffrey Hawkes, Stefan Bertilsson

## Abstract

Dissolved organic matter (DOM) is ubiquitous in aquatic ecosystems and fundamental for planetary processes and ecosystem functioning. While the link between microbial community composition and heterotrophic utilization of DOM has been recognized, the full diversity of organic compounds, their bioavailability, degradability and specific influences on the diversity and function of heterotrophs are still not clear. Here we experimentally investigate heterotrophic bacteria in thirty-three freshwater model communities. We identified 34 different heterotrophs growing in ambient lake DOM with taxonomic affiliations matching abundant freshwater bacterioplankton. We further describe 25 different heterotrophs growing in the phycosphere of *M. Aeruginosa* and 6 heterotrophs growing on the DOM produced by *M. Aeruginosa* with taxonomic affiliation in accord to phycosphere heterotrophs. In our experiment we observed that heterotrophs that live in the phycosphere remove more dissolved organic carbon than abundant freshwater heterotrophs. Moreover, phycosphere heterotrophs have bigger genomes than abundant lake bacteria. Altogether of the 4224 chemical features that were resolved by LC-MS, only 1229 were seen in all three treatments. None of the common/shared compounds were removed across all the model communities, suggesting contrasting niches of the studied taxa. Altogether our study highlights how each model community, with its unique taxonomic assemblages and organotroph functioning is upkept by the chemodiversity of DOM.

## Introduction

Dissolved organic matter (DOM) is ubiquitous in aquatic ecosystems and fundamental for planetary processes. DOM consists of a complex and variable mixture of compounds that serve as substrates for heterotrophic microorganisms (Azam 1998, Ferrer-Gonzalez et al 2021, Patriarca et al 2020b). As long-term carbon repository (∼660 Gt C), DOM holds 200 times more carbon than aquatic living biomass and is equal to the carbon storage in the atmosphere (Bar-On et al 2018, Falkowski et al 2000, Hansell et al 2009, Koch et al 2008). The bulk of DOM in continental waters is typically allochthonous (derived from terrestrial sources), chemically complex, and biologically recalcitrant (Amon and Benner 1996, Koehler et al 2012) while autochthonous DOM is produced in-situ and typically more bioavailable and therefore rapidly consumed in a way that prevents build-up (Maki et al 2009, Patriarca et al 2020b). Aquatic microorganisms play central roles both as DOM producers and consumers and also connect these dissolved substrates to larger organisms in the food web via the microbial loop (Azam 1998, Fenchel 2008). Despite the significance of microorganisms as mediators of biogeochemical processes in aquatic environments, their ecological niches in the turnover of the DOM pool are still unresolved. This is at least in part because aquatic microbial diversity is high while many abundant microbes remain uncultivated (Rodríguez-Gijón et al 2022), and also because of the high complexity and dynamic nature of aquatic DOM (Patriarca et al 2020b).

Earlier studies have demonstrated that phylogenetically diverse marine phytoplankton species release distinct arrays of organic compounds into their phycosphere (Becker et al 2014). This attracts and selects for distinct groups of heterotrophic bacteria (Ferrer-Gonzalez et al 2021, Sarmento and Gasol 2012). Accordingly, the composition and diversity of freshly produced and released autochthonous DOM influences microbial community composition and diversity, and those microorganisms that profit will in turn interactively drive DOM turnover in the oceans. Neveretheless, the most abundant marine bacteria (Giovannoni et al 2005) are not particularly abundant in the phycosphere (Buchan et al 2014, Seymour et al 2017) suggesting a more cryptic and multifaceted role of autochthonous DOM as a factor controlling microbial community structure. The same pattern is seen in freshwater lakes, where microbes that are numerically abundant in planktonic communities (Newton et al 2011) are typically not prominent or abundant in the phycosphere (Cai et al 2014, Eiler et al 2006) suggesting niche specialization with regards to DOM use, possibly related to the much larger allochthonous component in the combined DOM pool (Patriarca et al 2020b).

To experimentally uncover links between heterotrophs and their consumption of DOM, a few pioneering studies have relied on mass spectrometry paired with simplified model systems with a few identified keystone species of heterotrophs (Ferrer-Gonzalez et al 2021, Uchimiya et al 2021). Furthermore, a study using fourteen radiolabeled substrates and microautoradiography and fluorescence in situ hybridization (MAR-FISH) found niche partitioning with regards to organic substrate use for different groups of abundant lake bacteria (Salcher et al 2013). While bacterial community composition and substrate utilization patterns are tightly linked in aquatic ecosystems (Logue et al 2016), the detailed relationship between the complexity and reactivity of DOM and the functionally diverse heterotrophs is still not well understood.

With this study we advance the understanding of how specific heterotrophic bacteria act as central agents in the microbial loop. As models we use model communities developed from lake bacterioplankton under selection (i) in coculture with the cyanobacterium *Microcystis aeruginosa* (ii) growing on DOM produced from an axenic culture of *M. aeruginosa*, and (iii) growing in ambient lake DOM. After dilutions and serial transfer, cultivated model communities were established and characterized with a powerful combination of comparative metagenomics and high-resolution untargeted metabolite analysis. The model communities served as tractable model systems with reduced complexity (Garcia 2016) to assess the role of DOM quality in selecting for different microbial cohorts, the reactivity of different DOM compounds and how the heterotrophs differed with regards to organic compound utilization patterns.

## Results and discussion

### Microbial assemblages grew faster on freshly produced DOM than on lake DOM

In total, we inoculated 990 cultures, of which only 33 feature stable growth and could be upscaled (Table S1). Out of these 33 cultures, 5 originated from inoculation on spent media from axenic *M. aeruginosa* (M-DOM; initial DOC 6.2 mg/L), 21 cultures originated from filtered sterilized lakewater (L-DOM; initial DOC 10.7 mg/L) and 7 cultures were established as co-cultures with *M. aeruginosa* (M-co; DOC without heterotrophs 14.6 mg/L) (Figure 1). For growth characteristics, some general observations were made; (i) the M-DOM cultures reached the highest cell densities and also featured the highest growth rates (Figure 2E), (ii) M-co cultures reached similar cell densities as L-DOM cultures, (iii) among L-DOM and M-DOM communities, those that reached higher cell densities also grew faster than those that reached lower cell densities (Figure 2E). Finally, heterotrophs in M-co cultures used on average more DOC (dissolved organic carbon) per cell per day, while heterotrophs in M-DOM cultures used the least amount of DOC (Figure 2F). In our experiment we can observe how heterotrophs that live in the phycosphere and are not considered among the most abundant aquatic microorganisms, remove more dissolved organic carbon than abundant heterotrophs.

**Figure 1.**
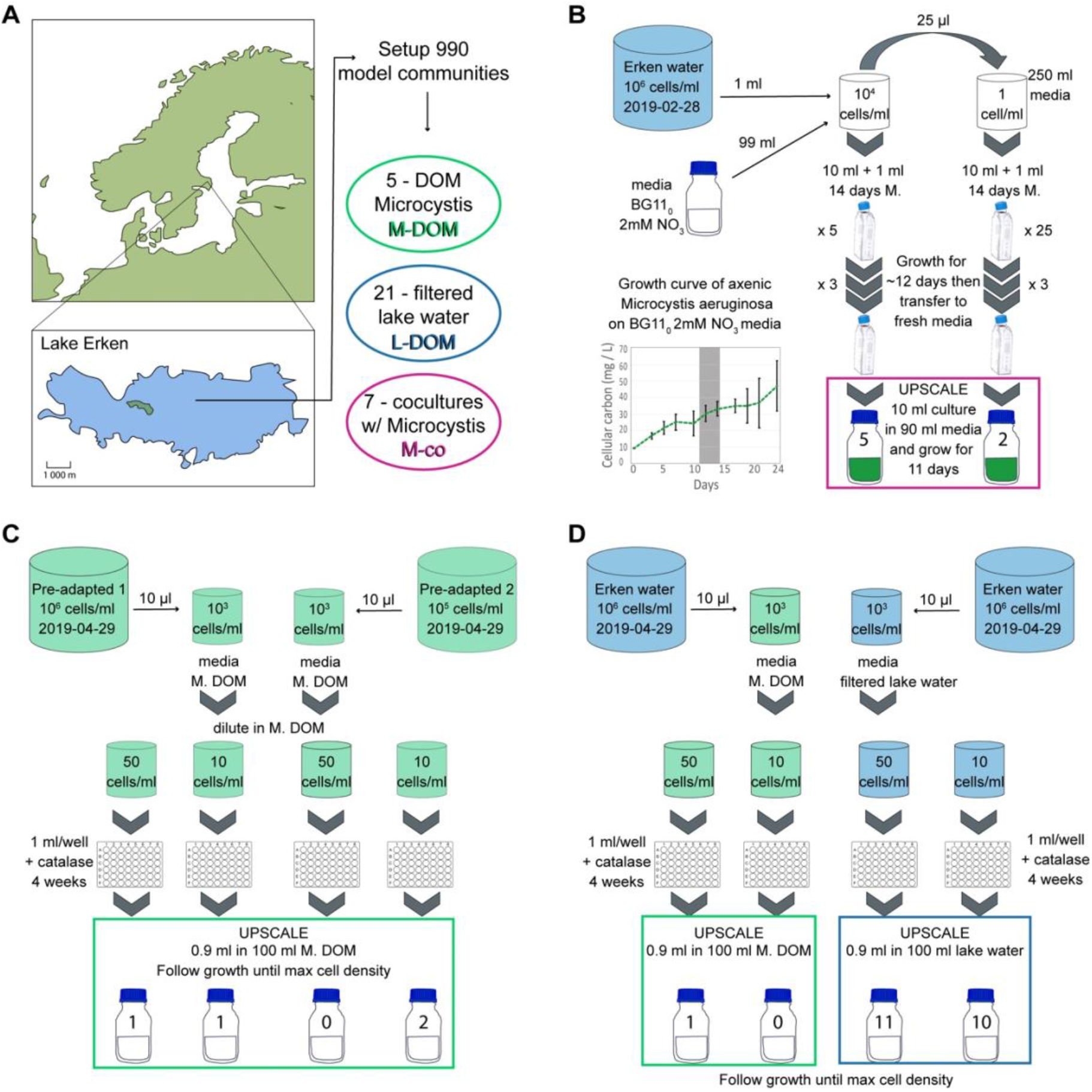
Overview of methods to establish model communities. Location of Lake Erken and overview of three treatments of model communities [A]. Seven model communities of Lake Erken cells grow together with Microcystis aeruginosa on BG11_0_ 2 mM NO_3_ media [B]. Pre-adaptation enrichment of cells from Lake Erken from February 2019 to M. aeruginosa DOM serve as inoculum for dilution cultivation of model communities in April 2019 yielding four model communities [C]. Cells from Lake Erken collected in April 2019 yield 21 model communities growing in Lake Erken DOM and 1 model community growing in DOM from M. aeruginosa [D].

**Figure 2.**
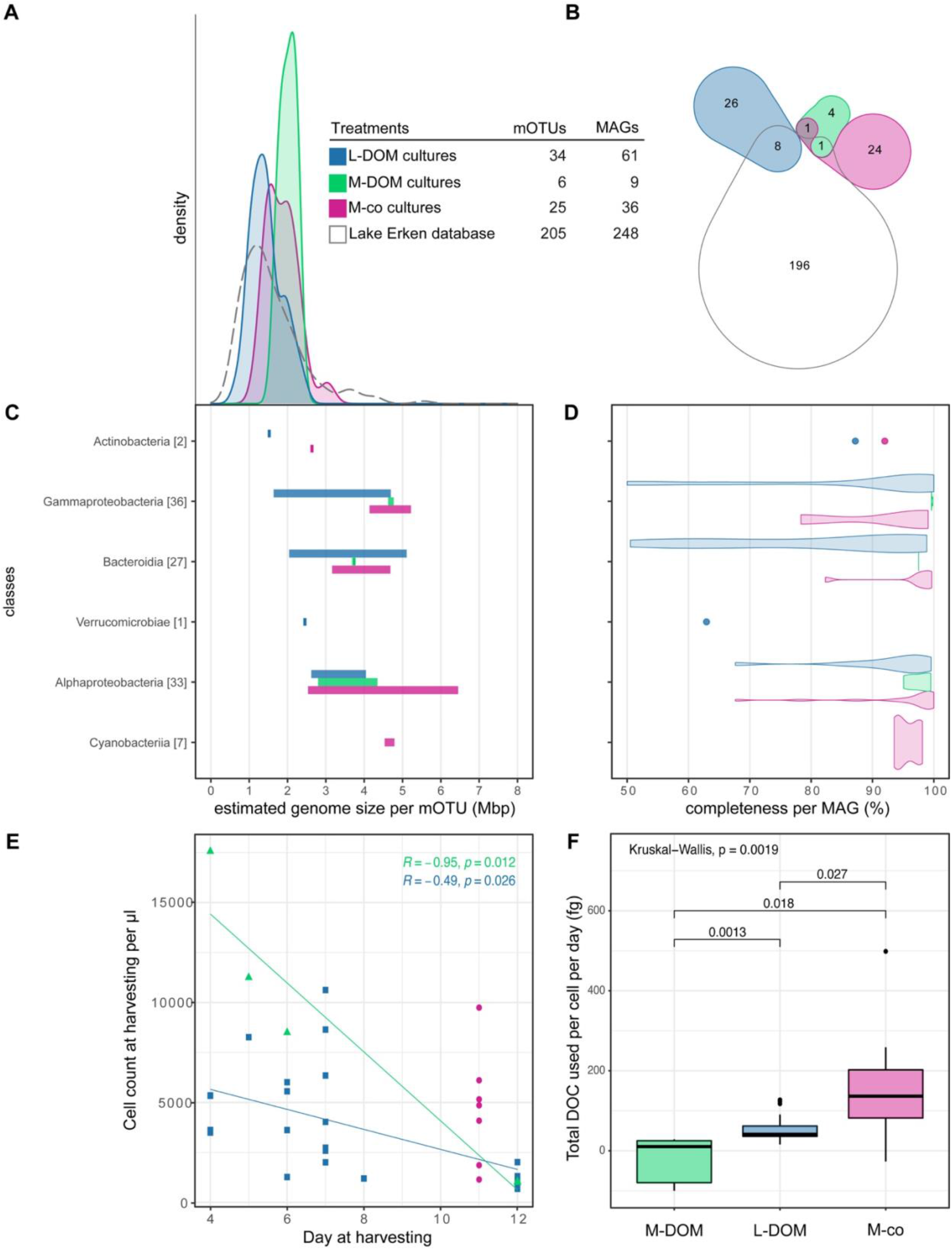
Overview of the genome size and growth across all model communities. Genome size distribution of bacteria in the 3 different treatments [A]. We use one representative metagenome assembled genome (MAG) per metagenomic operational taxonomic unit (mOTU - defined by 95% ANI). To construct the figures, we plotted the estimated genome sizes which were calculated based on the genome assembly size and completeness estimation provided. Grey dashed line includes 6251 MAGs which are representative genomes in the biggest freshwater genome collection until date. Venn diagram of the intersection between the representative MAGs of the three treatments [B]. The intersection was calculated using FastANI (Jain et al 2018) and was determined with a threshold of 95%. Genome size distribution across different bacterial classes [C]. Completeness distribution of the different MAGs calculated using CheckM (Parks et al 2015) [D]. Final day of growth, followed by harvesting vs. number of cells at harvesting per µl of each model community [E]. Cell count in model communities in coculture with M. aeruginosa includes count of M. aeruginosa. Total dissolved organic carbon (DOC) used per cell per day in mg per µl [F].

### Treatments are imprinted in the heterotroph average genome size

We reconstructed 61 metagenome-assembled genomes (MAGs) from the community assemblies in L-DOM cultures (Figure 2 and Table S1). These 61 MAGs grouped into 34 metagenomic operational taxonomic units (mOTUs) calculated based on 95% average nucleotide identity (ANI). 95% ANI is a widely recognized and used threshold for operationally defining the microbial equivalent to species (Garcia et al 2018b, Jain et al 2018). We further reconstructed 9 MAGs from M-DOM cultures that grouped into 6 mOTUs and an additional 36 MAGs from M-co cultures that grouped into 25 mOTUs. For context, we included a synoptic genomic inventory of the microbial source community in Lake Erken which included data from public databases (Buck et al 2021a, Mondav et al 2020). All 14 Lake Erken samples compiled yielded 248 MAGs that clustered in 205 mOTUs (Figure 2). The 354 MAGs were of good quality at >50% completeness and <5% contamination (Figure 2D) (Parks et al 2015).

We compiled and compared the genomes of representative MAGs from each mOTU (Table S3) for the different experimentally derived cultures as well as for Lake Erken. We found that the average estimated genome size was similar between M-DOM and M-co cultures (Wilcoxon, p=0.71) and MAGs from these two treatments had a significantly larger estimated genome size as compared to genomes from L-DOM cultures (Figure 2A and Figure S1, Wilcoxon p=0.019 and p=0.0001). The estimated genome size in the Lake Erken genomic survey encompassed the entire genome size range observed in the cultures (Figure 2A) but in average was smaller than genomes in M-DOM (Wilcoxon, p=0.07) and significantly smaller than those in M-co-cultures (Figure S1, Wilcoxon p=0.001). In conclusion, this implies that large genome size is characteristic for heterotrophs in the phycosphere. It is known that larger genomes require more maintenance (Giovannoni et al 2014), and the phycosphere provides an environment with labile and ‘plentiful’ resource.

Eight of the L-DOM treatment MAGs clustered in mOTUs with in-situ MAGs from Lake Erken (Figure 2B, MAGs marked in bold in Figure 3, Table S3 for details), while merely one of the MAGs from the M-DOM treatment clustered in an mOTU together with MAGs from Lake Erken. None of the MAGs from M-co cultures clustered together in mOTUs with in-situ MAGs from Lake Erken. Moreover, for most of the MAGs recovered from M-DOM and M-co culture treatment it was difficult to recruit reads from Lake Erken metagenomes that matched with more than 95% sequence identity (Figure 3). This suggests that microorganisms inhabiting the phycosphere are quantitatively scarce in community-level samples retrieved from the lake. The phycosphere is known as a hotspot for biogeochemical cycling (Seymour et al 2017) where a large proportion of the DOM released from phytoplankton cells are consumed (Smriga et al 2016).

**Figure 3.**
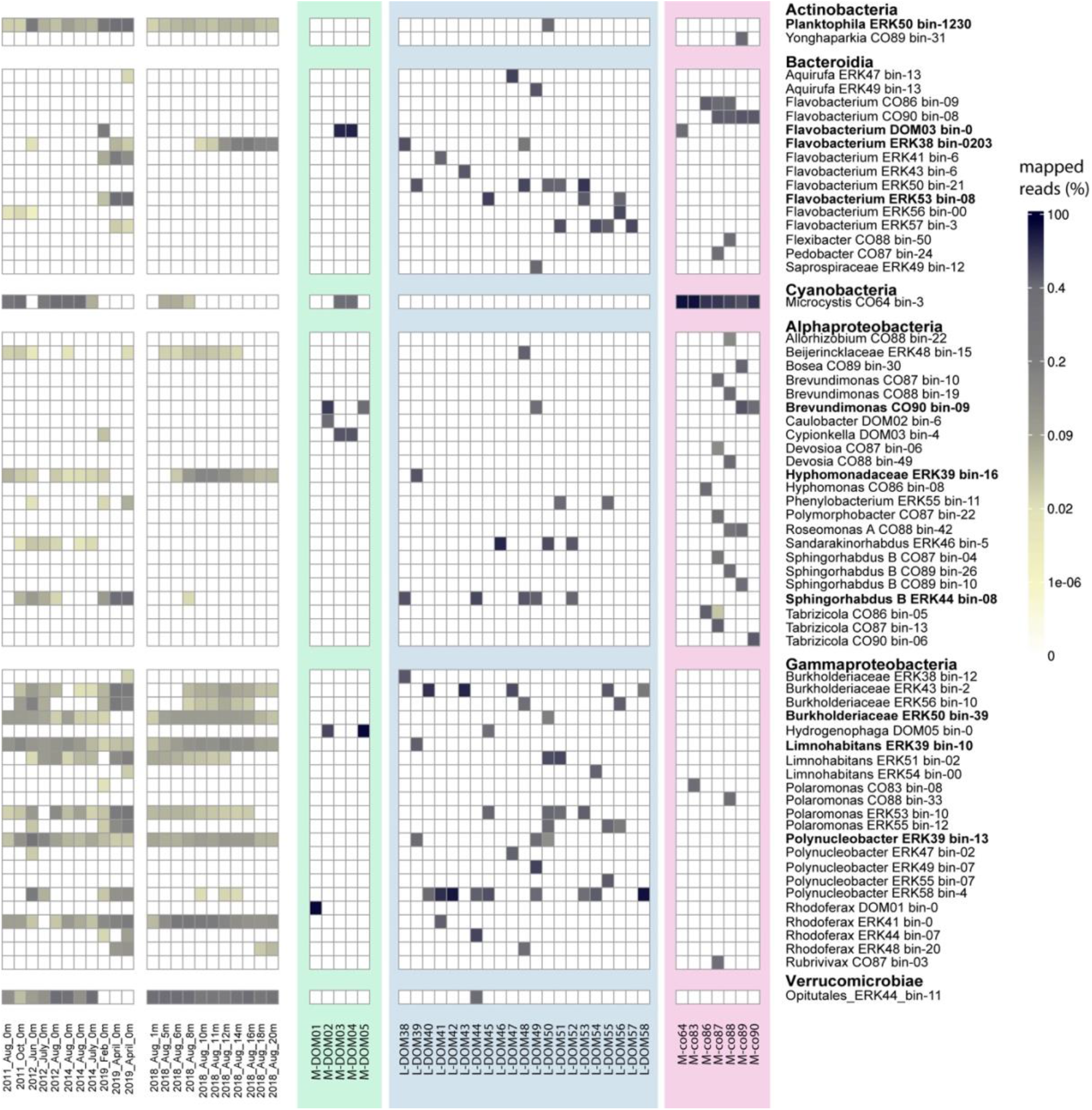
Relative read abundance for the 64 mOTU representatives of the 33 model communities and 22 Lake Erken metagenomes. The reads were normalized to the relative abundance of reads per metagenome. To show mapped reads, each representative MAG needed to have a breath coverage of 50%. Taxa are organized per Class and alphabetically. MAGs in bold are in the intersect between treatments or intersect with MAGs binned from the lake (Fig 2B).

### Taxonomic composition of bacterial diversity is treatment-dependent

All three treatments fostered communities with a distinct taxonomic signature. In the L-DOM cultures we retrieved lineages that are typical for freshwater lakes (Newton et al 2011), such as *Polynucleobacter* (Hahn et al 2016), *Planktophila* (Mondav et al 2020), *Caulobacterales* (Hentchel et al 2019) and *Limnohabitans* (Hahn et al 2010). In the M-co cultures, we recovered diverse *Alphaproteobacteria* affiliated with the orders *Sphingomonadales* and *Rhizobiales* as well as *Bacteroida* within the *Cytophagales* and *Flavobacterials*, most of them uncharacterized at the species level. The diversity was much lower in the M-DOM cultures, where recovered taxa represented *Hydrogenophaga, Caulobacter, Brevundimonas* and *Flavobacterium omnivorum*. A single mOTU included MAGs from both the M-DOM and M-co culture treatments (Figure2B, *Flavobacterium* DOM03 in bold in Figure 3). None of the mOTUs included MAGs from all three treatments. In our approach through dilution and serial transfers, we created reduced complexity sub-communities from the natural lake bacterioplankton assemblage. This approach enable diverse and abundant microorganisms to grow while maintaining interactions with partnering microorganisms that they also interact with in their natural environment (Garcia 2016, Pascual-Garcia et al 2020, Yu et al 2019). The observed taxonomic composition per treatment fit previous observations of abundant freshwater heterotrophs (Newton et al 2011, Salcher et al 2013) and the broader classes that are known to thrive on freshly produced phytoplankton DOM (Cai et al 2014, Eiler et al 2006, Ferrer-Gonzalez et al 2021).

Reconstructed good quality MAGs from cultures across all treatments grouped in 31 different genera. Amongst them, Flavobacterium with 22 representatives in 10 different mOTUs was the most prevalent. Flavobacterium representatives were recovered in 62% of the L-DOM cultures, 40% of M-DOM cultures, and 86% of the M-co cultures. The mOTUs of cultured Flavobacteria from L-DOM treatment were more abundant in the Lake Erken metagenomes collected in colder and deeper waters whereas the Flavobacterium from M-DOM and M-co cultures were hardly found in the lake (Figure 3). In contrast, only 2 additional Flavobacterium mOTUs were recovered from Lake Erken metagenomes and one of those was abundant in the warmer and upper water column (Figure S2). As the samples used to set up model communities of this study were collected in April 2019, their high prevalence in the emerging cultures is likely at least partly related to high abundances in the inoculum but may also be related to their previously recognized high capacity to exploit phytoplankton-derived organic matter (Williams et al 2013, Zeder et al 2009) and this is also supported by their metabolic profile suggesting a heterotrophic lifestyle.

*Polynucleobacter* is another prevalent genus with 11 representatives among reconstructed MAGs. These MAGs clustered in five different mOTUs and grew in 12 cultures, all of which were from the L-DOM treatment. These MAGs had an estimated genome size in the range of 2 to 2.6 Mb and GC content in the range of 43 to 49%. Metabolic reconstruction of these MAGs suggests a photoheterotrophic lifestyle with the capacity for aerobic anoxygenic phototrophy via pufM/L genes. An overall view of the metabolic potential of MAGs belonging to this genus also suggests thiosulfate oxidation via SOX complex.

### Genomes in different treatments vary in encoded functions

Comparative analysis of functional annotation of the MAGs recovered from each treatment show that more than 59% of all annotated functions (3516 out of 5918 unique KO identifiers) were shared among all three treatments (Venn diagram in Figure 4). These shared functions comprise 75.5, 89.2, and 66.3 percent, respectively, of the annotated functions in MAGs grown on L-DOM, Microcystis DOM, and M-co culture DOM treatments, and encode for core bacterial metabolic modules.

**Figure 4.**
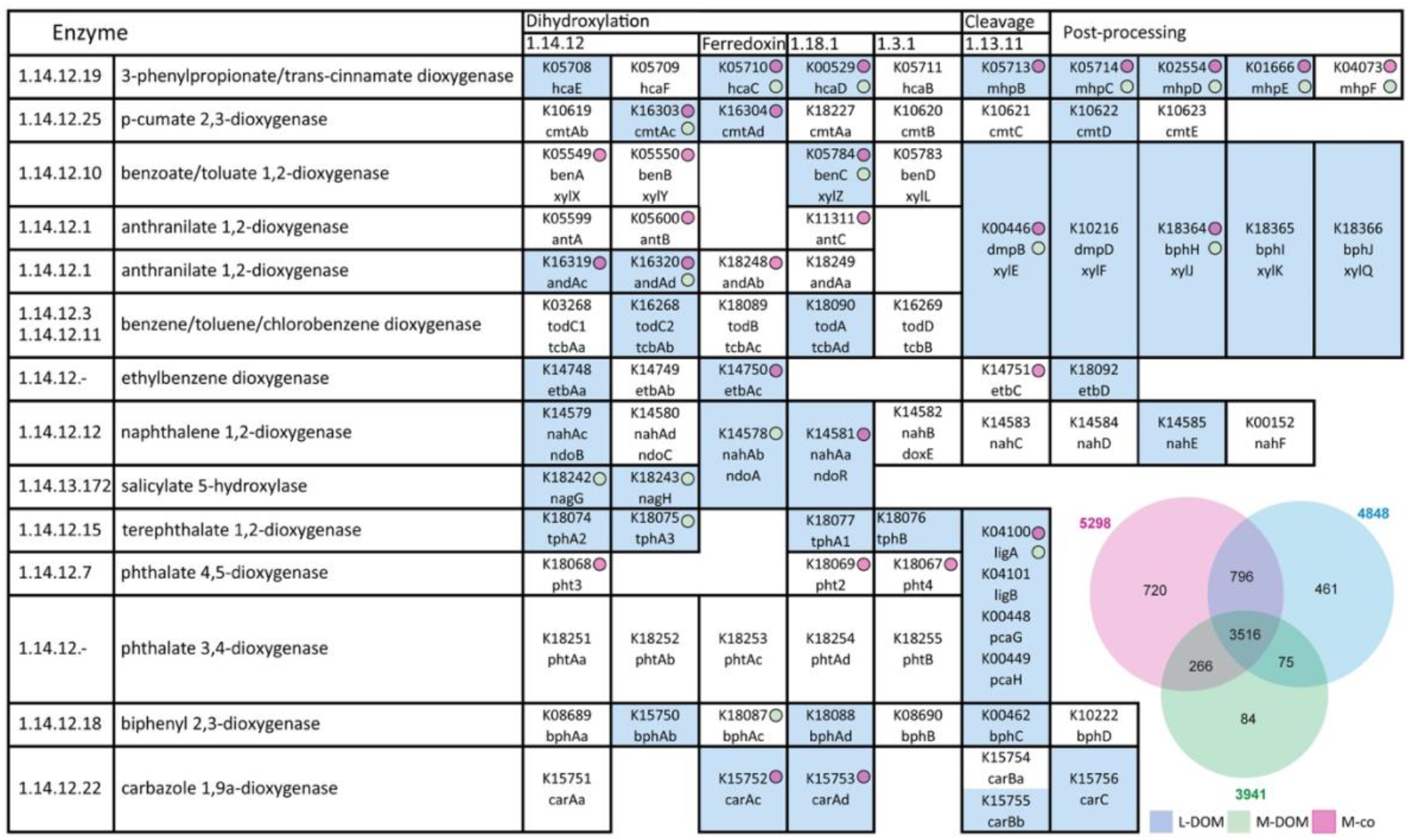
Distribution of dioxygenases involved in aromatic ring cleavage (KEGG brite br01602) in MAGs reconstructed from different treatments. Genes recovered from MAGs reconstructed form L-DOM cultures are highlighted in blue. Their presence in the other two treatments is also shown with circles. The Venn diagram shows the overlap of annotated functions in reconstructed MAGs from different treatments.

The 13.6% of annotated functions unique to MAGs reconstructed from M-co cultures encode for modules related to photosynthesis (found in the reconstructed Microcystis MAGs) and thiamine salvage pathway. Thiamin diphosphate that is the active form of thiamin, functions as the cofactor for transketolases, decarboxylases, and other enzymes that are involved in producing or breaking C-C bonds. At the genome level, this module (K00878 and K14153) is encoded in treatments M-co83 and M-co88 and in both cases in two representatives of the *Polaromonas* genus that are clustering in separate mOTUs. In our dataset, no other MAG was detected to encode for K00878 (hydroxyethylthiazole kinase), limiting this module to these two MAGs. This would point to thiamine salvage pathway being a possible candidate for community cooperation in the environment (Mondav et al 2020).

Functions shared between L-DOM and M-co cultures comprise 15% of total annotated functions in M-co cultures and encode for genes involved in degradation of aromatic compounds via a complete module of Catechol ortho-cleavage (K03381, K01856, K03464, K01055, and K14727). Catechol is a central intermediate in the degradation of different aromatic compounds and has previously been reported to also be produced in some algae (Singh et al 2017). At the level of individual genomes, we detected the full module in bin-30 MAG from the M-co89 treatment affiliated to genus *Bosea*, as well as bin-08 and bin-0 from L-DOM treatments taxonomically affiliated with Sphingomonadaceae and genus Ga0077559. Both of these MAGs cluster in mOTU_023 with two other MAGs from the L-DOM treatments (ERK38_bin-14 and ERK49_bin-02 with 67.6 and 97.3% completeness, respectively Table S2) where the latter two are missing genes involved in the Catechol ortho-cleavage module. This hints at a role of ecotype differentiation for DOM degradation potential (Mondav et al 2020). Interestingly members of mOTU_023 seem to grow with different microbial partners in the 4 different L-DOM model communities where they were present. Apart from the two Sphingomonadaceae MAGs, individual genes involved in Cathecol-ortho cleavage were also detected in other MAGs but not as a complete module.

The common annotated functions between M-co cultures and the M-DOM cultures (5%) encode for genes such as beta lactamase class D encoded in MAGs CO86_bin.08, affiliated to genus Hyphomonas and DOM03_bin-0 and DOM04_bin-10 clustering as mOTU_017 affiliated to genus Flavobacterium. Interestingly both of these Flavobacterium MAGs were reconstructed from M-DOM treatments that only contain one other MAG affiliated to Pseudorhodobacter_A (both of these also cluster in mOTU_150). The Pseudorhodobacter_A MAGs have photoheterotrophic metabolism with aerobic anoxygenic phototrophy via pufM/L genes. The Pseudorhodobacter_A MAGs also encode complete modules for biosynthesis of cobalamin and can synthetize cobyrinate a,c-diamide via the aerobic pathway as opposed to Microcystis that encode the anaerobic pathway for cobyrinate a,c-diamide production via sirohydrochlorin.

Among genes uniquely retrieved in the L-DOM treatments (9.5% of all annotated functions in these treatments) we observed several different Dioxygenases. One of the most important functions of dioxygenases is the cleavage of aromatic rings and they are widely found in nature (Harayama and Rekik 1989). L-DOM MAGs only share a few functions with the M-DOM MAGs, but this include some dioxygenases suggested to be involved in naphthalene degradation (Figure 4).

Genome-encoded functions unique to the M-DOM treatments (only 2% of the annotated functions in this treatments) encode for beta lactamases and biphenyl 2,3-dioxygenase ferredoxin component. The highest diversity of dioxygenases was detected in the L-DOM cultures. Apart from the fact that most of the reconstructed MAGs belong to the L-DOM cultures (n=61) compared to M-co cultures (n=36) and M-DOM cultures (n=9), this higher diversity could also be due to higher representation of aromatic compounds in the native DOM pool of the L-DOM treatment as compared to the cyanobacterial DOM treatments (Patriarca et al 2020b).

### Chemodiversity of aquatic DOM sustain bacterial diversity

We discuss our data in terms of chemical features, which are MS/MS signals. One metabolite can lead to several features, so we refrain from naming them metabolites or compounds. Altogether 4224 LC-MS features were resolved and had sufficient peak intensity to trigger data-dependent MS/MS analysis in at least one sample in the dataset. Of these features, 3417 were present in at least one sample above 5x the intensity of the average of 4 blanks. Depending on the culture type (M-DOM, L-DOM or M-co cultures), a subset of these 3417 features were detected in the initial feed (see first row of Figure 5B). The M-DOM, L-DOM and M-co feed had 2422, 1872 and 2146 features respectively, with only a minor fraction unique to any individual feed (Figure 5C). 1229 features were found in all three feeds. Variable numbers of features were either removed or produced in the incubation replicates, leading to presence/absence and intensity differences in the sample extracts (Figure 5B). Pairwise sample dissimilarities were calculated with the Bray-Curtis metric, allowing samples to be compared and grouped in a principal coordinate diagram (PCoA) (Figure 5A). The L-DOM incubation samples were clearly separated from the M-DOM and M-co cultures along the first principal coordinate, while the latter two groups were separated along the second, indicating that the treatments varied more than the variance among biological replicates within each group. A PERMANOVA analysis of the Bray-Curtis distance matrix found an F-statistic of 22.1 for group membership (L-DOM, M-DOM or M-co), with a p value of 0.01 over 100 iterations of the PERMANOVA algorithm. As expected, the controls for M-DOM (medium incubated without heterotrophs) and the M-co cultures (*M. aeruginosa* cultivated without heterotrophs) were similar while they diverged along PCoA2 during the experimental incubations.

**Figure 5.**
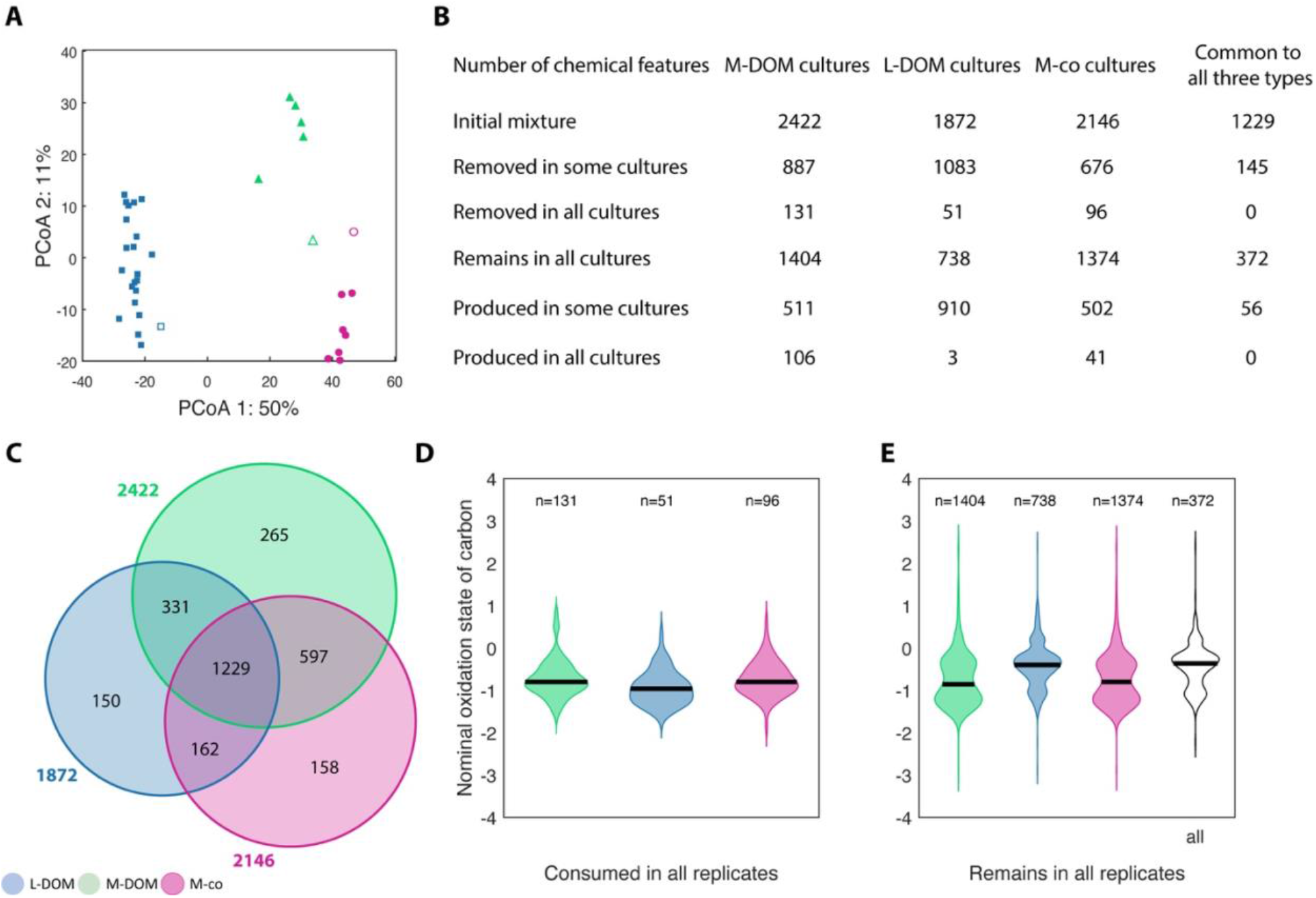
PCoA of the chemical composition of DOM in the model communities [A]. Filled symbols are each of the cultures, whereas empty symbols are the media control without heterotrophs. Table of number of metabolites in the initial mixture, removed in some or all cultures, remaining in all cultures or produced in some or all cultures [B]. Venn Diagram with the number of initial chemical features in the different treatments [C]. Violin plot of the oxidation state in the consumed metabolites [D]. Violin plot of the oxidation state of the metabolites that remain [E].

Of the 1229 features found in all three feeds, 372 remained in all incubations thoruhgout the experimental incubation. None of the common/shared feed features were removed across all of the cultures, indicating niche partitioning on substrate consumption by the taxa cultivated in the different treatments. However, for each feed type, there was a moderate number of features (between 51 and 131) that were removed in every culture with this experimental group (see third row of Figure 5B). We suggest that these features represent the most broadly bioavailable substrates in each treatment. Taken together, the presence/absence results indicate that variability between the different cultures and treatments was high and that each model community, with its unique taxa and functions is upkept by the chemodiversity of DOM.

We observed that heterotrophs in M-DOM cultures consumed more lipids, organic acids and organoheterocyclic compounds, whereas M-co cultures consumed more organic oxygen compounds (Table S4). Taxa in L-DOM cultures consumed less compounds that could be resolved and classified with the methods used and may rely more on the unresolved compounds typical for allochthonous DOM (Figure 2F and Table S4). We further calculated the nominal oxidation state of carbon (NOSC) for the features that were consumed and those that remained in all biological replicates for the respective treatment (Figure 5D-E). The mean and range of values was rather similar between feed types for consumed features but differed for the features that remained in all cultures between M-DOM/M-co cultures and the L-DOM cultures. The remaining features L-DOM cultures were substantially more oxidized than those remaining in the M-DOM or M-co cultures.

### Relative oxidation state of labile and stable features

We expected most of the highly bio-labile features to be more reduced, and more stable features to be more oxidized, as more energy should be available from oxidation of reduced species (Chen et al 2021). The only treatment where this pattern was found was in L-DOM, where the features consumed in all replicates vs. stable in all replicates had a median NOSC of -0.96 and -0.4 respectively (Figure 5D-E). Generally speaking, and particularly in the treatments fed cyanobacterial DOM or co-cultures with Microcystis, the NOSC varied widely with many features having values -1.5 to 0, with higher variance in stable features compared with those consumed. This implies a diversity of metabolites with differing functionalities both in the bio-labile and stable groups.

### Limitations and outlook

Here we report on heterotroph use of DOM in inoculated cultures amended either with cyanobacterial metabolites or the natural organic matter of lake water. The observed LCMS features derive from dissolved organic compounds in the samples that are either present in the initial feeds, produced by autotrophs or resulting from biomolecular transformation processes during the experimental incubations. Such DOM mixtures are extremely complex, and resolvable features such as those reported in the present study only represent a fraction of the total DOM. Additionally, the more labile low molecular metabolites autotrophically produced during the incubations may not be detected in chemical assays such as the LCMS analysis used here, simply because of their rapid assimilation and use while they can still be among the most important heterotrophic substrates.

Additionally, the bulk of DOM in freshwater lakes is unresolvable alicyclic carboxylic acids and lignin derived polymers that have limited value as heterotrophic substrates and originate from incomplete breakdown and molecular transformation of terrestrial organic matter in soils (Hertkorn et al 2006, Patriarca et al 2020a, Zherebker et al 2017). We found that many organic matter features in our dataset were resistant to removal across the relatively short timescale of our experiment, but this does not exclude the possibility that such recalcitrant DOM may degrades slowly over many months to years in natural inland waters (Koehler et al 2012, Mostovaya et al 2016). In conclusion, our approach is useful to compare metabolites in the experiment with DOM found in nature because the fragmentation pattern is like a signature that would otherwise be hidden in molecular formulas.

Cultivation experiments are known to bias against cultivation of many abundant microorganisms (Lewis et al 2020, Swan et al 2013). We used a dilution-to-extinction mixed cultivation approach (Garcia 2016) and obtained some abundant microorganisms in culture. However, the cultivation success was poorer than expected. One explanation could be that all our experimental incubations were done at 20°C and while this is optimal for *M. aeruginosa* growth, it is substantially higher that in situ conditions in the lake which was 4°C in February and 10°C in April. This explains why although our cultures hosted some of the most abundant freshwater bacterial groups, there was less success in establishing viable cultures as compared to similar experiments where incubation temperature was close to *in situ* conditions (Garcia et al 2018a). Earlier work has shown that freshwater bacteria feature microdiversity with variable temperature optima (Garcia-Garcia et al 2019). It is possible that at the time of sample collection cells that were more abundant in-situ were more viable at 4°C or 10°C.

Despite these limitations, our study was able to retrieve relevant microbial subsets and characterize these with metagenomics and high-resolution mass spectrometry to find the different microbes living and feeding of different types of DOM. We found that the phycosphere heterotrophs consume more dissolved organic carbon as well as had bigger genome sizes. In general, the bigger genome sizes in heterotrophs consuming autochthonous DOM allowed them to encode for cobalamin biosynthesis pathway and thiamine salvage pathway, which are a function rare in aquatic environments (Garcia et al 2015, Giovannoni 2012, Paerl et al 2018). This points to dependencies between heterotrophs of different kinds. Moreover, we different genes encoding for degradation of aromatic compounds, dioxygenases or beta lactamase in different heterotrophs feeding on lake DOM and primary produced DOM. And while the chemodiversity of DOM holds a niche for many different heterotrophs, we found in our study that these heterotrophs might not only depend on these chemodiversity, but also help upkeep it with their production. In our study, we confirm the high carbon consumption of heterotrophs in the phycosphere and speculate that their turnover in the microbial loop might also be high, since phycosphere microbes found to be abundant in aquatic ecosystems. Finally, in our study we cultivated thirty-three model communities, but we believe that high-throughput cultivation, sequencing and analytical chemical approaches will further reveal the roles of the diversity of microorganisms in aquatic environments.

## Methods

### Environmental sampling and DNA extraction

We collected environmental samples from the surface layer of Lake Erken on February 4 °C and April 2019 10 °C and DOC 10.7 mg/L. Lake water samples were collected both for cultivation and to create a background genomic library (Supor-100 0.1 µm). Samples were transported to the laboratory within one hour of collection and further processed by 0.1 µm membrane filtration (Hollow fiber cartridge, GE healthcare) to produce media for the L-DOM cultivations. Filters from lake water samples and harvested upscaled cultures were used to extract DNA using the DNasy PowerWater kit (Qiagen) following the quick-start protocol.

### Cultivation of model communities

All cells growing in model communities were collected from Lake Erken in February or April 2019. To obtain simplified sub-communities we used dilution methods in 3 types of media: 0.1 µm filtered Lake Erken water, 0.1 µm filtrate of *Microcystis aeruginosa* that grew on BG11_0_ media (Rippka et al 1979) or BG11_0_ medium amended with actively growing *Microcyctis* cells (Figure 1). The cyanobacterial cells were provided from the Pasteur Culture Collections of Cyanobacteria (PCC, France). All cultures were grown with a photon flux density of 25 μmol m−2 s−1 (Philips TL-D 18&30W 865 Super 80) under a 12-hour light/dark cycle at 20 °C. We have two main types of cultivation setups (i) dilution of Lake Erken cells in order to cultivate between 10 and 50 cells in 1 ml medium in a 96 well plate either in 0.1 µm filtrate from Lake Erken (L-DOM cultures) or 0.22 µm filtrate from axenic Microcystis cultures (M-DOM cultures) (Figure 1C-D), (ii) dilution of Lake Erken cells in order to grow between 10 and 10 000 cells together with about 10^7^ Microcystis cells for four 10% inoculum transfers in fresh BG11_0_ media (M-co cultures) (Figure 1 B). We set up dilution to extinction experiments in Feb 2019 but had no growth in the wells. However, the pre-dilutions performed in February on 0.22 µm filtrate from axenic Microcystis cultures were incubated at 20 °C and tested for cell count in April. One of the pre-dilutions had 10^6^ cells per mL and was considered pre-adapted inoculum one used to set up model communities in April (Figure 1C). Another of the pre-dilutions showed 10^5^ cells per mL and was considered a pre-adapted inoculum two.

The L-DOM and M-DOM cultures (Figure 1C-D) were screened for growth and selected for upscaling if cell densities were greater than 1000 cells per µl after 4 weeks incubation in 96-well plates. After selection of 50 such cultures for upscaling, these were grown in 100 ml of their respective media in Schott bottles under a 12-hour light/dark cycle at 20 °C. The upscaled cultures were monitored for growth by daily aseptic subsampling and subsequent flow cytometry cell counts (CytoFLEX, Beckman Coulter). Cultures were harvested when cell densities did not change during two consecutive days and was close to the cell density of the culture used to inoculate the upscaling. Cells were harvested by vacuum filtration onto a Supor-100 0.1 µm membrane filter (Pall, USA). M-co cultures were transferred three times about 12 days into cultivation under a 12-hour light/dark cycle at 20 °C in 10 ml volume flasks to ensure the selection of actively growing cells (Figure 1 B). M-co cultures were upscaled to 100 ml for the fourth transfer and harvested in 0.22 µm Sterivex filters (Millipore) on day 11 of the incubation.

### Flow cytometry

All cell counts were performed in a CytoFLEX (Beckman Coulter). Acquisition settings, FSC 2500, SSC 2500, FITC 800, PC5.5 257. Primary threshold FITC Automatic. Flow rate 60 µl/min. Cells were stained using Syto13 (Invitrogen). Original Syto13 solution at 5 mM in DMSO. A Syto13 working solution was prepared from the original solution by diluting 1 to 200 in milliq water. Then 50 µl of sample were stained with 2.6 µl of working solution of Syto13.

### Library preparations and sequencing

DNA samples were prepared in 1/10 of a volume of the standard library protocol using the Mosquito HV liquid handler from SPT Labtech (formerly TTP Labtech, Melbourn, UK). Libraries were prepared from extracted DNA of metagenomic samples in the range of 0.1 - 201 ng/µl using the Nextera Flex kit now known as Illumina DNA Prep kit (Illumina, San Diego, CA). Using the Mosquito HV (SPT Labtech, Melbourn, UK), reaction volumes were scaled down to a 1/10 of a volume compared to the standard Illumina protocol with the following exceptions: the post-tagmentation washing buffer (TWB) was added in a volume 8 ul (2 × 4 µl) and the 80% ethanol during library clean-up was added in a volume of 4,5 µl. Three µl of DNA diluted to an input of approximate 3-10 ng/µl was added to a premixed solution of bead linked transposons (BLT) and tagmentation buffer (TB1).

A 384-bottom plate magnet (SPT Labtech, Melbourn, UK) was used for all steps involving binding and washing of magnetic beads. Tagmented DNA samples with an input of 10-24 ng/µl were run using 8 library amplification cycles and those with 1-9 ng/µl at 12 cycles using unique dual index combinations, IDT for Illumina DNA/RNA UD Indexes set A (Illumina, San Diego, CA) to mitigate index hopping. After double sided library clean-up using AMPure XP beads (Beckman Coulter, Brea, CA) libraries were resuspended in 10 µl, of which three µl was aspirated and diluted in nine µl, to account for quality control and quantification. A subset of 20 samples were run on a bioanalyzer or fragment analyser, consistently showing an average total fragment length of 500-600 bp. Each library was quantified using the KAPA Library Quantification kit (ROCHE, Basel, CH) in three dilutions per sample for calculation of molarity and dilution before equimolar pooling of 1 nM each. Due to the low molarity in the total volume of both pools, they were dried down prior to sequencing. The library pools were sequenced on a NovaSeq 6000 SP flow cell, PE 2 × 150 bp with 10% PhiX.

### Metagenome assemblies and postprocessing

The raw data was first trimmed using Trimmomatic (version 0.36; parameters: ILLUMINACLIP:TruSeq3-PE.fa:2:30:10 LEADING:3 TRAILING:3 SLIDINGWINDOW:4:15 MINLEN:36) (Bolger et al 2014). The trimmed data was assembled using Megahit (version 1.1.13) (Li et al 2015) with default settings. The relevant quality controlled reads were mapped to all the assemblies using BBmap (Bushnell 2016) with default settings and the mapping results were used to bin the contigs using Metabat (version 2.12.1, parameters --maxP 93 --minS 50 -m 1500 -s 10000) (Kang et al 2015). Genes of obtained MAGs were predicted using Prokka (version 1.13.3, default settings) (Seemann 2014) and annotated using eggNOG-mapper (version 2.0.1, default settings) (Cantalapiedra et al 2021, Huerta-Cepas et al 2019). Prokaryotic completeness and redundancy of all bins from Metabat and for all assembled single cells were computed using CheckM (version 1.0.13) (Parks et al 2015). The MAGs were clustered into metagenomic Operational Taxonomic Units (mOTUs) using mOTUlizer version (0.2.2) (Buck et al 2021b) starting with 50% complete genomes with less than 5% contamination. Genome pairs with ANI 95% or more were clustered into connected components. Additionally, less complete genomes were recruited to the mOTU if its ANI similarity was above 95%. MAGs were taxonomically annotated using GTDB-Tk (version 102 with database release 89) (Parks et al 2018). Mapping for the heatmap was done with Bowtie 2 for relative abundance estimates in the samples (‘--ignore-quals --mp 1,1 --np 1 --rdg 0,1 --rfg 0,1 --score-min L,0,-0.05’) (version 2.3.5.1) (Langmead and Salzberg 2012). MAG coverage with breadth threshold of >= 50% was used to confirm a true positive.

### Liquid Chromatography Tandem Mass Spectrometry analysis

Metabolites were analyzed in the samples after solid phase extraction on a hydrophobic sorbent at acidic pH. The extracted samples were separated and analyzed in duplicate using ultrahigh performance liquid chromatography coupled to positive mode electrospray ionization high resolution mass spectrometry (Orbitrap Q-Exactive). Analytes emerging from the column were fragmented using a data dependent analysis routine, allowing for feature matching to libraries of previously reported compounds and putative formula assignment, as detailed below. Solid phase extraction was used to de-salt and concentrate analytes. Cartridges (Agilent PPL; 3ml, 100 mg) were rinsed with one column volume of methanol and 0.1% formic acid. Samples (∼5 ml), which were stored frozen in Falcon tubes, were thawed and acidified with 50µL 10% formic acid containing 300ppb Capsaicin as an internal standard. The samples were loaded by gravity onto the cartridges, which were then rinsed with a column volume of 0.1% formic acid and dried by vacuum. The analytes were eluted with 1.3ml methanol by gravity, the methanol was dried and the samples redissolved in 100 µL of 5:95 LCMS grade acetonitrile:water with 0.1% formic acid, containing 100 ppb Hippuric acid and Fusidic acid and 1 ppm Raffinose, as internal standards.

Samples were injected at 10 µL into the UPLC system, and were separated on a Phenomenex Kinetex C18 column (2×150 mm, 1.7 µm) at a flow rate of 0.4 ml/min. Mobile phase A was 0.1% formic acid in LCMS grade water, B was 0.15% formic acid in LCMS grade acetonitrile. A linear two step gradient started at 5 % B then started to increase at 30 sec from 5-50% B at 7 min followed by an increase to 99% B at 10 min, a 3 min washout phase at 99% B, and a 4 min equilibration phase at 5% B (method length = 17 min). MS1 resolution was set to 70000, MS2 resolution 17500. Data dependent analysis was set to be activated for the top 5 peaks, with a trigger range of 2-15 s and dynamic exclusion of 5 s, 1 Da isolation width and stepped collision energy at 20,30,40 eV. mzXML files were generated from Thermo .raw files with ReAdW.

Molecular formulas for the resulting MS2 features were generated using SIRIUS and ZODIAC software (Duhrkop et al 2019, Ludwig et al 2020). SIRIUS uses isotope pattern matching and MS/MS fragmentation trees (Bocker and Rasche 2008) to rank possible molecular formulas. ZODIAC uses molecular formulas generated via SIRIUS and re-ranks them based on the MS/MS network topology to each other. We used 10 ppm as default mass accuracy for both MS1 and MS/MS level and C, H, N, O, P, S as possible elements for de novo structure elucidation in addition to the molecular formulas in the biodatabases in SIRIUS. The nominal oxidation state of carbon (NOSC) was calculated as in Equation 1:

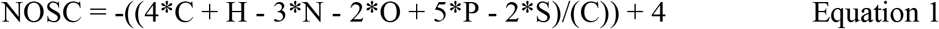

The minimum and maximum values for NOSC are -4 (e.g. CH4) and +4 (e.g. CO2).

Feature intensities were normalized per sample and used for sample dissimilarity calculation via the Bray-Curtis metric, which is common in ecology. The Bray Curtis dissimilarity matrix was subsequently used as the basis for a principal coordinate analysis via classical multidimensional scaling (cmdscale, MATLAB version 2017b).

Peak abundances are highly influenced by sample contents at the ESI spray, which can cause suppression or promotion, and may drift during a long analytical run. For this reason, four internal standards were added to the extracts (one before SPE (Capsaicin) and three after (Hippuric acid, Fusidic acid and Raffinose)) in order to assess the suitability of the method for comparing feature abundances between samples. The standards were always detected, but the relative standard deviation of these four standards over 36 samples and 4 blanks (after duplicate averaging) was 40, 66, 24 and 26%, respectively. Due to this relatively large variability, which is likely due to variable suppression effects, we only consider features that were removed below or emerged above the detection limit when we mention metabolite consumption or production.

## Data availability

All raw sequencing data is available under ERP124195, PRJEB37497 and PRJEB38681, and all accession numbers are found in Table S1. All raw mass spectrometry data is available through the MassIVE repository (massive.ucsd.edu) under the following accession number: MSV000086759.

## Acknowledgements

We are grateful to John Paul Balmonte for helpful discussions. We thank Kamila Koprowska, application scientist at SPT Labtech for helpful technical assistance setting up the Mosquito HV protocol.

The work was primarily funded by Science for Life Laboratory, Knut and Alice Wallenberg Foundations (grant KAW 2013.0091), Kungl. Vetenskapsakademiens stiftelser (CR2019-0060) and the Swedish Research Council (grant 2017-04422). The authors would like to acknowledge support from the Genomics infrastructure services at Science for Life Laboratory (https://www.scilifelab.se) in Uppsala. The metagenomics libraries were prepared at the Microbial Single Cell facility and sequencing was performed by the SNP&SEQ Technology Platform in Uppsala. The facilities are part of the National Genomics Infrastructure (NGI) Sweden and Science for Life Laboratory. The SNP&SEQ Platform is also supported by the Swedish Research Council and the Knut and Alice Wallenberg Foundation. The computations and data handling were enabled by resources in the project SNIC 2021/6-99 and SNIC 2021/5-133 provided by the Swedish National Infrastructure for Computing (SNIC) at UPPMAX, partially funded by the Swedish Research Council through grant agreement no. 2018-05973. We thank the Deutsche Forschungsgemeinschaft for the support of D.P. through the CMFI Cluster of Excellence (EXC 2124).

## Author contribution

SLG, VTSN, MB and SB conceptualized the research idea. SLG and VTSN performed all cultivation experiments. SLG, JH, AMD, MB, JN, MM, and DP performed data analysis. SLG drafted the first manuscript. JH, MM and JJH wrote some sections of the manuscript. All authors contributed to writing and editing of the manuscript.

## Figures and Supplementary Material

**Figure S1.**
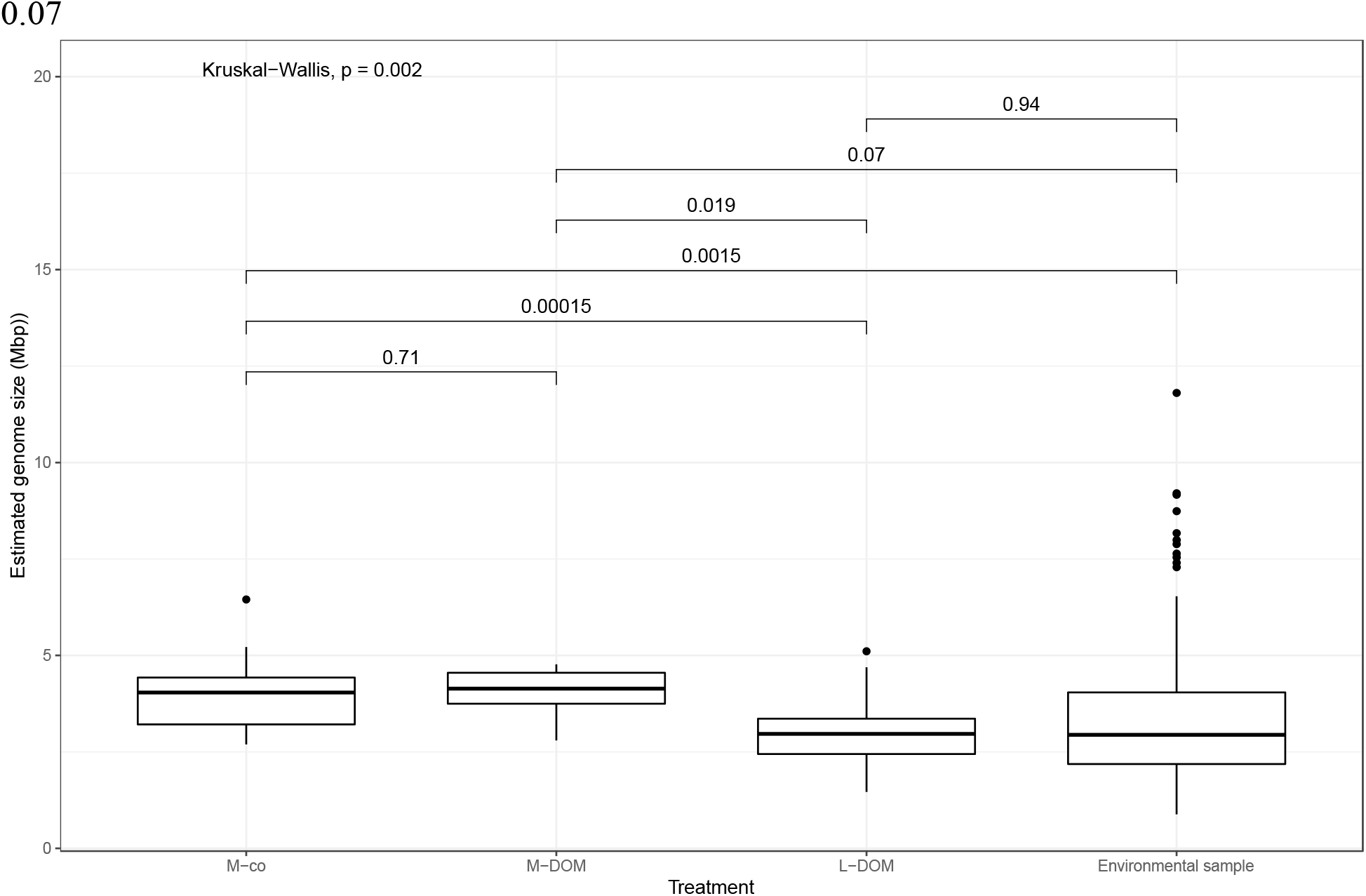
Average estimated genome size per treatment and Wilcoxon statistical comparison.

**Figure S2.**
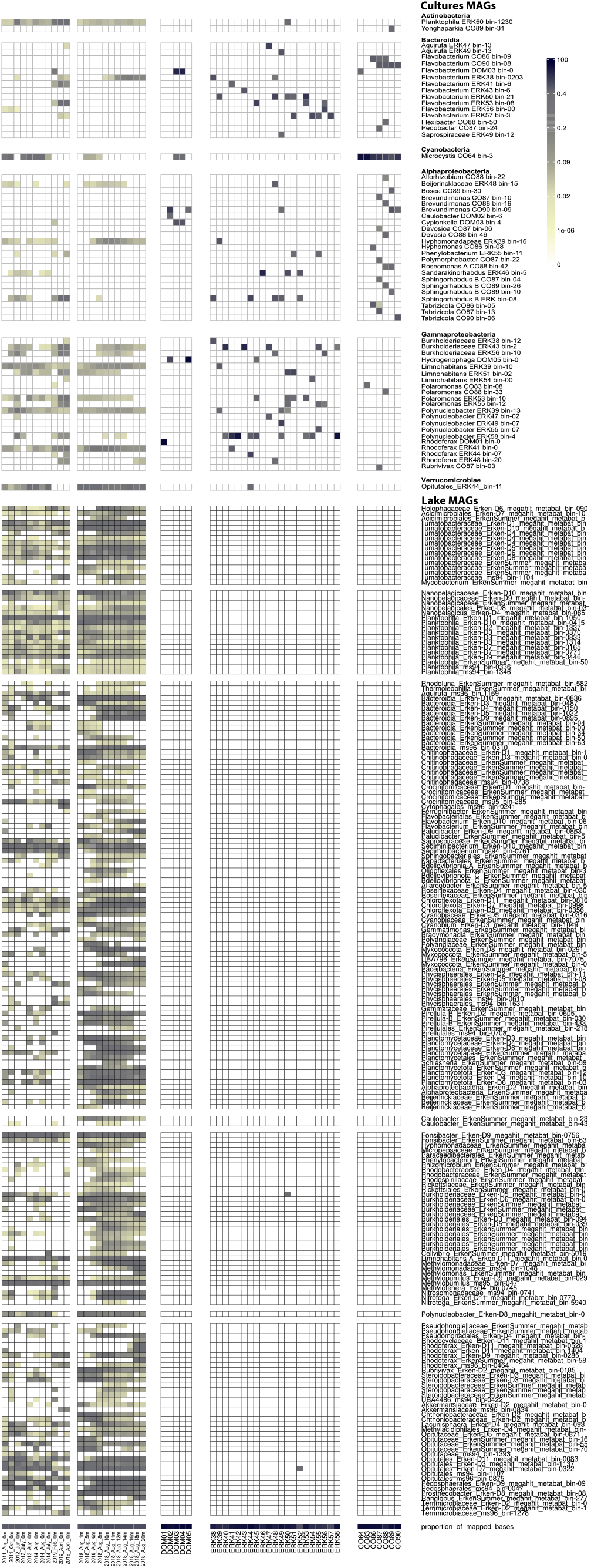
Relative read abundance for the 260 mOTU representatives of the 33 model communities and 22 Lake Erken metagenomes. The reads were normalized to the relative abundance of reads per metagenome. To show mapped reads, each representative MAG needed to have a breath coverage of 50%. Taxa are organized per Class and alphabetically.

Table S1. Accession numbers and metadata of 33 model community metagenomes and 3 Lake Erken metagenomes

Table S2. Information about all MAGs binned from the 33 model community metagenomes

Table S3. Information about all MAGs that were selected as representatives for the mOTUs in the study

Table S4. Compounds types removed in all cultures per treatment.

